# Behavioral and metabolic effects of extended hippocampal overexpression of the myokine Irisin in an Alzheimer’s Disease mouse model

**DOI:** 10.1101/2024.12.16.628747

**Authors:** Jacob A. Borukhin, Erin Kang, Rodrigo F. Tomas, Caoyuanhui Wang, Arielle Trotz, Adrian Requejo, Jeremy C. McIntyre, Karina Alviña

**Affiliations:** Department of Neuroscience, University of Florida, Gainesville, FL 32610; McKnight Brain Institute, University of Florida, Gainesville, FL 32610. Department of Neuroscience, University of Florida, Gainesville, FL 32610

## Abstract

Lifestyle interventions such as exercise are promising strategies to reduce the detrimental effects faced by aging populations. In particular, the exercise-induced myokine Irisin could act as a mediator of the positive effects of exercise, even as a potential therapeutic option for Alzheimer’s disease (AD). However, the fundamental roles of Irisin in counteracting AD neuropathology and other effects have not been fully determined. In addition, whether centrally produced Irisin has any effects is unclear. Here we examined the neurobehavioral effects of overexpressing Irisin directly in the hippocampus of an AD transgenic mouse line (TgCRND8), using adult male mice stereotaxically injected with Irisin-AAV vector. To further mimic chronic exercise, spatial memory and exploratory behaviors were assessed using the novel object recognition (NOR)/open field test (OFT) and Y-Maze tests after 8 weeks.

Our results showed that Irisin expression led to several behavioral changes, including increased grooming behavior and slightly improved spatial memory performance in short interval NOR test in Tg mice. In the Y-Maze test however, we observed an increase in time spent in familiar spaces, which could indicate heightened anxiety-like behaviors in mice injected with Irisin-AAV. In addition, we observed significant differences in body and liver size, and in circulating metabolites such as ketone bodies, that were differentially regulated in NTg or Tg mice.

Overall, our findings show important neurobehavioral effects of long-lasting overexpression of the exerkine Irisin in the hippocampus. Our results also raise questions about potential detrimental effects and thus warrant further studies to fully dissect how this exercise mediator affects the brain.

## Introduction

Exercise has powerful and well-established physiological effects on different systems in the human body, including the central nervous system. Examples of such effects include promoting neurogenesis, neuronal plasticity and survival ^1,2^, which have been shown to help age-related neurodegenerative diseases. These effects can also offer a non-pharmacological approach to remedying the cognitive decline associated with aging ^3^. Of these neurodegenerative diseases, Alzheimer’s Disease (AD) remains at the forefront of therapeutic research, as this disease has impacted more than 6.9 million citizens aged 65 and older, and this number is only expanding without effective treatment for the associated physiological and psychological deficits. Because of its known positive effects, mechanisms of neuroprotection associated with exercise need to be fully understood and further elucidated. While robust evidence has provided insight into the molecular basis of exercise-induced neuroprotection, many biomarkers of exercise, and their relation to neuroprotection, remain underexplored, ultimately limiting the potential of developing novel therapeutics against neurodegenerative diseases like AD.

Fibronectin type-III domain-containing protein-5 (FNDC5) is a transmembrane protein expressed in several tissues including skeletal muscles. When stimulated by exercise, FNDC5 is enzymatically cleaved to form a soluble secretory peptide called Irisin ^4^. Irisin is a peroxisome-proliferator activated receptor gamma (PPARγ)-coactivator 1α (PGC-1α)-dependent myokine that is capable of browning adipose tissue in both mice and humans, effectively promoting thermogenesis during exercise ^4^. While FNDC5 is primarily found within skeletal muscles, it has also been localized to the brain, specifically within the hippocampus, a region heavily involved in memory and cognition. Within the hippocampus, increased expression of FNDC5 mRNA in response to exercise has been shown to stimulate the expression of Brain Derived Neurotropic Factor (BDNF) ^5^, a neurotrophin that plays a vital role in neurogenesis, synapse formation and plasticity, learning, and memory. These findings suggest that exercise has the capacity to enhance neuroprotection through a FNDC5/Irisin/BDNF mechanism ^5,6^. Such paradigm raises the potential for FNDC5/Irisin to act as a novel therapeutic in combating many age-related neurodegenerative disorders, namely AD and associated dementias.

In the present study, we aimed at testing the potential of brain derived Irisin to attenuate the behavioral deficits associated with AD using a well-established mouse model. By combining adeno-associated viral (AAV) vector technology to drive the expression of Irisin in the hippocampus with behavioral tests, we tested the hypothesis that overexpression of Irisin in the hippocampus might rescue memory deficits in male AD mice. Simultaneously, we assessed exploratory and anxiety-like behaviors, and overall metabolic status of injected mice. Our results show that overexpression of Irisin in the hippocampus has the potential to prevent memory impairment associated with AD, albeit at the cost of potentially increasing anxiety-like behaviors. Overall, our results provide additional support toward developing a novel therapeutic for AD mediated by the myokine/exerkine Irisin.

## Methods

All procedures detailed here were approved by the University of Florida (UF) IACUC (Animal Protocol #202011308) and follow the U.S. National Institutes of Health (NIH) guidelines for animal research.

### Experimental Animals

Adult 12–15-month-old non-transgenic (NTg, control, n=14) and transgenic (Tg, n=12) CRND8 male mice were obtained from a UF colony and bred in house using B6C3F1/J female breeders obtained from Jackson Laboratories. Litters were genotyped, separated by sex and housed (maximum 5 mice per cage) in ventilated cages inside an air-conditioned room. The room was kept at 24°C and on a 12:12h *inverted* light/dark cycle (off 7:00am/ on 7:00pm). Mice had ad libitum access to standard laboratory food and water. In all experiments, each mouse was considered as an independent unit and tested only once. All behavioral experiments were performed in a separate room inside the animal facility where mice were kept.

TgCRND8 mice carry a double-mutant form of amyloid precursor protein (APP) 695 (K670N/M671L and V717F). These mice thus display an age-related increase in APP and Aβ peptide production and plaque deposition ^7^. TgCRND8 mice exhibit progressive plaque pathology beginning at 3 months of age. By 7 months, they display hyperphosphorylation and nitrosylation of tau along with cholinergic cell loss ^8^ TgCRND8 mice exhibited greater spontaneous behavior activity compared to wild type mice ^9^.

### Generation of Adeno associated virus (AAV) vector

To develop the Irisin overexpressing vector, standard cloning techniques were used. Specifically, PstI -NotI DNA fragments containing Irisin encoding sequence were generated using the pCR-blunt-TOPO-FNDC5 plasmid (Addgene: #35970), and the following primers were used for High Fidelity PCR: Irisin 5’ PstI: CAGTGTGATGGATATCTGCAGA, Irisin 3’ NotI: CATGCGGCCGCCTCCTTCATGGTCAC.

The PCR products and the pENN.AAV.CB7.CI.PM20D1-flag-WPRE-rBG (Addgene: #132682) containing a 6x-his tag were digested with PstI and NotI. The digested DNA was purified using gel and re-ligated using T4 DNA Ligase (NEB, Cat# M0202S). Lastly, the construct was sent to UF’s Ocular Gene Therapy Core facility for packaging in an AAV9 stereotype.

### Protein expression quantification

After completing behavioral experiments, hippocampal tissue was dissected and flash frozen for later processing. To measure the effect of AAV injection on protein expression, hippocampal tissue was processed to extract protein, and samples were then separated using standard western blot protocols. Specifically, dissected tissue samples were homogenized in RIPA buffer (150mM NaCl, 50mM Tris-HCl, 1% Nonidet P-40, 0.5% sodium deoxycholate, 0.1% SDS, and 5mM EDTA with Proteinase/Phosphatase inhibitors), centrifuged for total protein extraction. Extracted protein was quantified using commercial BCA protein assay kit (Thermo Scientific Cat# 23227) and stored at −80 °C until use.

Protein samples were mixed with Laemmli lysis buffer (Bio-Rad Cat#: 1610747), incubated for 5 min at 95°C and then separated using sodium dodecyl sulfate-polyacrylamide gel electrophoresis, on 4-20% Mini-PROTEAN TGX Precast Gel (Bio-Rad Cat# 456-8095). The separated proteins were then transferred onto nitrocellulose membrane using a Trans Blot Mini Cell (Bio-Rad Cat# 1703930) at 100V for 1h. For immunodetection, the blots were blocked with 5% dry milk in TBST (TBS+0.1% Tween-20) for 1h at room temperature, then incubated with primary antibodies overnight, at 4°C. Then blots were washed 3x in TBST and incubated with horseradish peroxidase (HRP) conjugated anti-rabbit or anti-mouse antibody (1:2000, MilliporeSigma Cat# AP307P, Invitrogen Cat#31430) for 1h at room temperature. After three washes in TBST, the bands were detected using Pierce™ ECL 2 Western Blotting Substrate kit (Thermo Scientific Cat# 80197). The images were captured with a Bio-Rad ChemiDoc MP Imaging System, and immunoreactive bands were quantified with Image Lab Software (Bio-Rad). The following primary antibodies were used: anti-irisin (1:1000, Phoenix Pharmaceuticals Cat# G-067-29), and anti-β-actin (1:1000, Sigma-Aldrich Cat# A1978).

### Stereotaxic Surgery

TgCRND8 and NTgCRND8 mice were randomly assigned to a different experimental group: injection with an adeno-associated viral (AAV) vector containing a scrambled (non-coding) sequence or injection with an AAV vector containing an Irisin encoding sequence. For stereotaxic surgery, mice were anesthetized with isoflurane and placed on a stereotaxic frame (Kopf Instruments). The skull was exposed, two small holes were drilled using a microdrill (Stoelting, Inc.) and the following coordinates from the Mouse Brain Atlas were used to target the dentate gyrus (DG) region of the hippocampus: anteroposterior (AP) 2.0mm, mediolateral (ML) 1.8mm, and dorsoventral (DV) 2.2mm. The injection was controlled via a micropump (World Precision Instruments, Inc.) connected to a Hamilton syringe that delivered 1-1.5 µL of the AAV solution. After the injection, the syringe was left in place for 5min, slowly withdrawn, and then the incision was closed with sterile sutures and biological tissue adhesive. After surgery, mice were transferred to a clean empty cage, placed on a heat pad for 45 min to 1h while being monitored for recovery. After completion of this period, mice were returned to their home cages and kept for up to 8 weeks to allow for the expression of the Irisin vector and to mimic chronic exercise paradigms. After this period, the first session of OF/NOR testing began, followed by Y-Maze testing.

### Behavioral Analysis

#### Combined Novel Object Recognition (NOR) and Open Field (OF) Test

We used a combined open field (OF) and novel object recognition (NOR) test that proceeded in three sessions: one habituation phase (OF/NOR1) followed by two recognition phases at different intervals: OF/NOR2 (1h after OF/NOR1) and OF/NOR3 (5min after OF/NOR2) for all mice (see schematic in Figure 1A). In the OF/NOR1 (habituation phase), we placed individual mice in a square OF arena (plastic container, 18” per side) for 10min while videotaping the session for behavioral analysis. Two identical objects (“A”) were placed in the arena and mice were allowed to freely explore and interact with. As objects for NOR we used one of the following: small 100 ml glass bottles wrapped with pink laboratory adhesive tape, a green plastic pumpkin, a blue water bottle and a red mason jar wrapped with red laboratory tape. The objects were all similar in height (∼4 in.), but differed in structure, texture and color. After the 10min OF/NOR1 (habituation phase) mice were removed from the OF arena and placed back in their home cages for 1h. After this 1h period, mice were brought back to the testing arena and were subjected to OF/NOR2 (recognition phase 1). For this test, one object of the identical pair was replaced with a novel object (“B”), chosen from the three different objects not used for habituation. Mice were placed back into the OF arena and allowed to explore for 5min while videotaping the session for behavioral analysis. After this 5min period of exploration, mice were again removed from the OF arena and placed back into their home cages for 5min. After this 5min period, mice were finally subject to OF/NOR3 (recognition phase 2). For this final test “B” was replaced with a final novel object (“C”), chosen from the two remaining objects not used for habituation or OF/NOR2. The same procedure from OF/NOR2 was followed for OF/NOR3. Short term memory (STM) function of the mice was quantified by analyzing exploratory behaviors during both recognition tests (OF/NOR2 and OF/NOR3).

**Figure 1.**
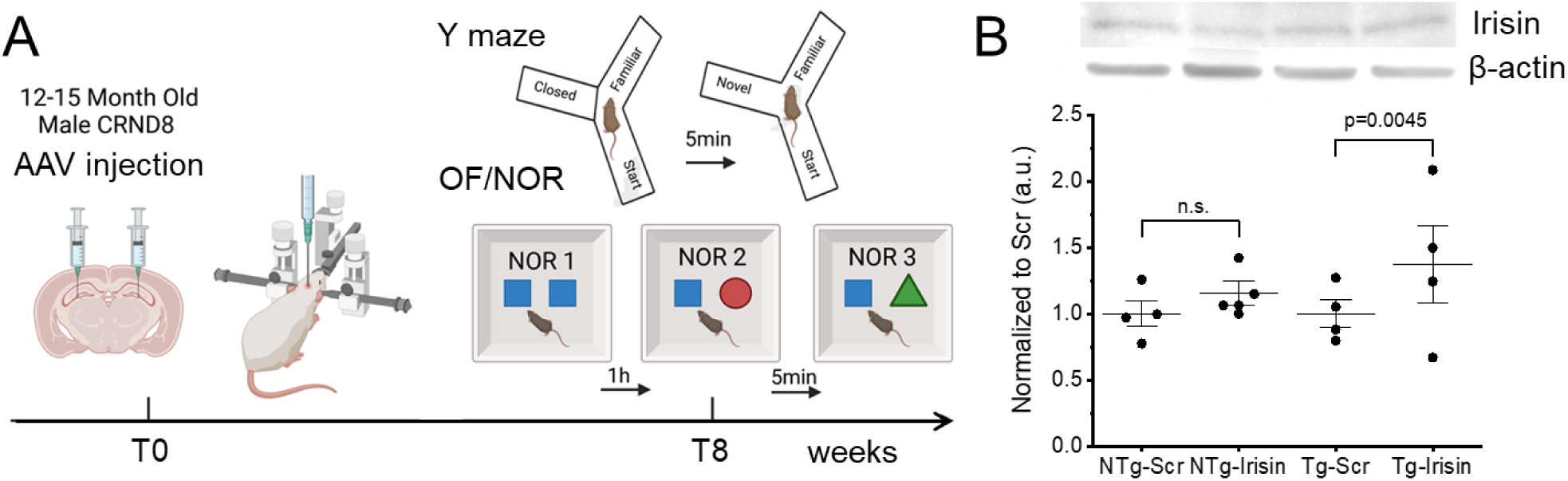
Schematic of the experimental timeline used in this study (A). Mice were stereotaxically injected with AAV-Irisin or scrambled (Scr, controls) at time zero (T0). 8 weeks post injection (T8), behavioral analysis was conducted using OF/NOR and Y maze tests. (B) Representative image of western blot and quantification of Irisin protein expression in the hippocampus of injected mice (n=4 per group). Statistical significance was evaluated by a Two-sample T test for each genotype, N.s. denotes not significant.

Exploratory behavior was defined as mice directly interacting with either object (i.e., sniffing, rearing on it, or directing the nose toward the object), analyzed using EthoVision XT Software Version 10.1 (Noldus Information Technologies, Inc.). Data from mice that did not meet the minimum exploration threshold of 1s exploration time on either object were not included. A “preference index” (PI) was then calculated for both OF/NOR2 and OF/NOR3 using the following formula: PI(OF/NOR2) = [Time on B / (Time on B + Time on A)] and PI(OF/NOR3) = [Time on C/ (Time on C+ Time on A)]. We then calculated the change in PI from OF/NOR2 to OF/NOR3 as ΔNOR using the formula ΔNOR = PI(OF/NOR3)-PI(OF/NOR2).

Locomotor and anxiety-like behaviors were also assessed during both the habituation and recognition phases. For analysis, the OF arena was divided into two equally sized areas comprising the center and periphery. Time spent in the center was quantified using analysis software (EthoVision XT Version 10.1, Noldus Information Technologies, Inc.). Trained observers blind to the treatment also manually analyzed rearing events (both supported and unsupported) and grooming time.

#### Y Maze Test

A Y-shaped maze apparatus (Stoelting Inc.) was used to test short term and spatial working memory in mice. On day 1, mice were initially placed in the lower arm of the Y-Maze and were allowed to acclimate and freely explore for 10 min. Mice were placed back into their home cage until day 2. On day 2, Y-Maze testing followed a similar pattern to OF/NOR testing with an initial habituation phase followed by a recognition phase. In the habituation phase, mice were placed at the edge of the middle arm of the Y-Maze (labeled as the “Start” arm), and one of the side arms was blocked, limiting exploration to two arms. Mice were allowed to explore the “Start” and the open arm (“Familiar”) for 10 min. Following the habituation session, mice were removed from the Y-Maze and were placed back into their home cages for 5 min. Then, the closed arm was unblocked and opened for the mice to explore (“Novel” arm). After the 5 min rest period, mice were placed back into the same “Start” arm of the Y maze and were allowed to freely explore all 3 arms (“Start,” “Familiar,” and “Novel”) while videotaping for behavioral analysis. Short term and spatial working memory function were quantified by analyzing exploration (time spent and distance covered) in all 3 arms during the recognition phase of the Y-Maze test. Time spent in each arm and distance traveled were quantified using video tracking and analysis software (EthoVision XT Version 10.1, Noldus Information Technologies, Inc.).

#### Body Weight Monitoring, Tissue Collection and Analysis

Body weight (BW) was monitored by weighing each mouse on a benchtop scale before stereotaxic surgery (before condition) and then before behavioral tests were conducted (after condition). Weight change was then calculated using the formula ΔBW = [(BW_before_ — BW_after_) / (BW_before_)] x 100.

Following completion of all behavioral testing, mice were anesthetized by isoflurane inhalation and euthanized via decapitation. Trunk blood was collected in sterile microcentrifuge tubes and then centrifuged for 15 min at 6000 RPM. The serum (supernatant) was then collected, aliquoted and placed in storage at –80°F for further analysis. In addition, several organs were dissected and weighed, including brain, liver, thymus, kidneys, subcutaneous adipose tissue, gonadal adipose tissue, perirenal adipose tissue, and mesenteric adipose tissue. To account for individual BW differences that could explain differences in organ size, organ weight values were normalized by each mouse BW value. All adipose tissue weights were summed into a total adipose tissue category and was used in determining the adiposity index (AI) of each mice using the formula AI = [Total Adipose tissue weight/BW_afte_r].

### Data Analysis and Sharing

All statistical analyses were performed by Origin software (OriginLab Corporation). We used a Two-way analysis of variance (ANOVA) to determine statistical significance between groups, and Tukey’s mean comparison post-test, or Two-sample T-test, as specified in each case. For all results, a level of p<0.05 was considered statistically significant. All raw and processed data are available upon request.

## Results

### Stereotaxic injection of AAV-Irisin vector results in increased expression of Irisin in the hippocampus of Tg mice but not in NTg counterparts

To test the effects of Irisin overexpression in the hippocampus of CRND8 AD mice we stereotaxically injected an AAV9-Irisin vector directly into the DG of Tg and NTg counterparts. As controls, we used a similar AAV carrying a scrambled sequence (Scr). Mice recovered from the surgery and were kept for up to 8 weeks to mimic the conditions of chronic exercise. After this period, behavioral tests were conducted, and mice were then euthanized to collect several tissues for further analysis (Figure 1A for timeline).

Hippocampal tissue was processed for protein extraction and then separated by electrophoresis (SDS-PAGE). The expression of Irisin was probed by Western blotting, as shown in the representative blot in Figure 1B. Quantification (and normalization using β-actin) showed that the expression of Irisin was higher in samples from Tg mice injected with AAV-Irisin compared to tissue from Tg mice injected with Scr vector (Figure 1B), however this difference was not observed in samples from NTg mice. On average, we found an approximately 16% increase in expression in NTg mice injected with AAV-Irisin (1.00±0.09 in NTg-Scr vs 1.16±0.09 in NTg-Irisin, n=4, p=0.289, Two-sample T-test). This increase rose to ∼37% in Tg mice injected with AAV-Irisin (1.00±0.11 in Tg-Irisin vs 1.37±0.29 in in Tg-Irisin, n=4, p=0.0045, Two-sample T-test).

These results show that our viral approach was effective in increasing expression of Irisin centrally, specifically in the hippocampus. It is interesting that this increase was significant in Tg mice only. This might be related to the fact that lower levels of Irisin have been previously found in AD mouse lines ^10^.

### Baseline Behavior during OF/NOR1 test

To establish baseline behavior in all treatment groups, a combined OF/NOR1 paradigm was used, and OF parameters were studied. Figure 2A shows that Tg mice moved more in the OF compared to their NTg counterparts, regardless of treatment. Tg-Scr and Tg-Irisin mice traveled on average 5583±1277cm (n=7) and 4362±1495cm (n=5), respectively, in the OF test compared to NTg-Scr and NTg-Irisin mice which traveled on average 2449±252.9cm (n=7) and 1602±265.2cm (n=7), respectively in the OF test (p=0.03 between genotypes, and not significant between AAV types, p>0.05). To further support this result, heat maps of each mouse were generated and were analyzed to assess the degree of movement in the OF/NOR1 test (Figure 2B). Comparing the representative heat maps, it is evident that NTg mice spent less time moving around the OF test, independent of treatment. Tg mice, on the other hand, traveled more as indicated by the higher heat density in the periphery. Additional analysis showed a trend towards less time in the center of the OF in Tg mice injected with AAV-Irisin (Figure 2C), although this difference did not reach statistical significance, possibly due to large variability within groups. On average, Tg-Scr mice spent 116.4±20.23s in the center, compared to Tg-Irisin mice which spent 62.71±23.30s in the center. Similarly, NTg-Scr mice spent 65.34±13.71s in the center vs NTg-Irisin mice that spent 53.70±20.23s in the center (p>0.05 for genotype and treatment).

**Figure 2.**
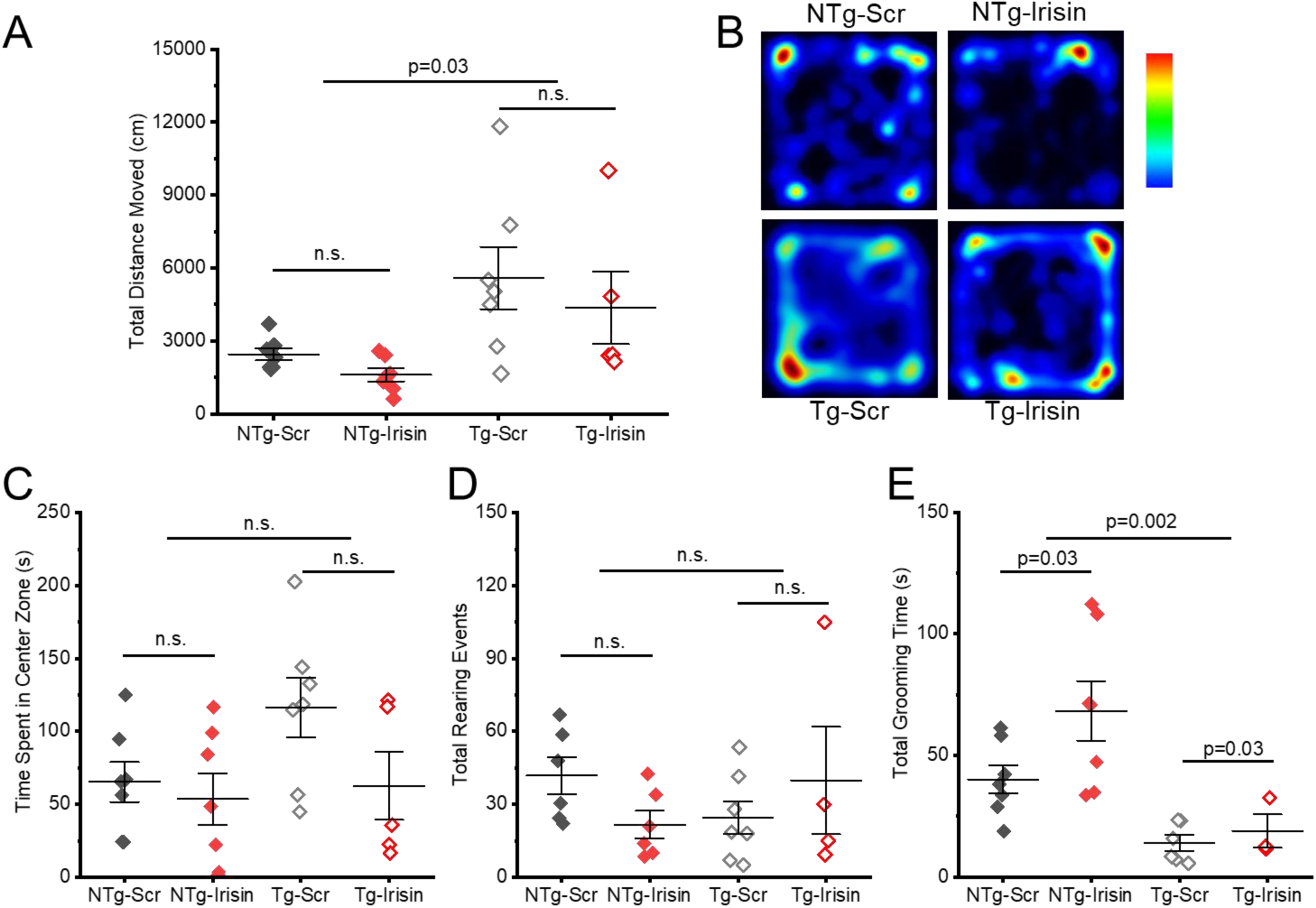
Baseline behavior during the habituation session of OF/NOR test (OF/NOR1). Panel A shows that Tg mice moved more than NTg counterparts, independent of injected AAV. (B) shows examples of individual mice during OF/NOR1 depicted as heat maps. No differences were observed in the time spent in the center of the arena (C) or total rearing events (D). Grooming behavior was significantly higher in mice injected with AAV-Irisin, regardless of genotype (E). N.s. denotes not significant, p>0.05, Two-way ANOVA.

To further explore anxiety-like and exploratory behaviors in our combined OF/NOR1 test, we manually assessed grooming time and rearing events (both supported and unsupported) in all experimental groups. Exploration of mice was gauged by monitoring two types of rearing events: “supported” rearing (i.e., when mice leaned on the walls of the OF arena), and “unsupported” (i.e., when they stood up on their hind legs). While there was a clear trend to less rearing events in Tg mice and because of AAV-Irisin injection, these differences did not reach statistical significance. NTg-Scr mice had an average number of rearing events of 41.78±7.70 events, which was more than NTg-Irisin mice, which had on average 21.67±5.64 rearing events; Tg-Scr mice had an average number of rearing events of 24.5±6.74 events, which was more than Tg-Irisin mice, which had on average 18.11±6.16 rearing events (Figure 2D, genotype and treatment, p>0.05). Furthermore, when analyzing grooming behavior, we found that both NTg and Tg mice spent significantly more time grooming when injected with AAV-Irisin (Figure 2E). On average, NTg-Scr mice (n=7) spent 40.15±5.78s grooming compared to NTg-Irisin mice (n=7), which spent 68.29±12.26s grooming (p=0.03); Tg-Scr mice spent 13.89±3.28s (n=6) grooming compared to Tg-Irisin mice, which spent 18.89±6.85s grooming (p=0.03, n=3). When comparing genotype effect, there was a significant difference (p=0.002) between NTg and Tg mice.

Taken together, these results show that TgCRND8 mice, regardless of treatment with Irisin, display higher locomotor activity in the OF test compared to NTg counterparts, consistent with previous findings ^11^. Additionally, an interesting increase in anxiety-like behaviors was seen in all mice injected with AAV-irisin, in agreement with previous findings ^12^.

### Effects of AAV-Irisin on Short-Term Memory in TgCRND8 Mice

Systemic Irisin has been shown to enhance working memory in AD mouse models and has the capability to prevent the cognitive defects associated with this disease ^13,14^. However, these previous examples have used peripheral Irisin, not exclusively centrally produced. To test the potentially restorative effects of Irisin directly overexpressed in the hippocampus, we used a combined open field (OF) and novel object recognition (NOR) test that we had previously characterized ^15^ and is well established in assessing short-term memory of mouse models ^16^. During the habituation phase (combined OF/NOR1), all mice familiarized themselves with the OF/NOR arena, and two of the same objects. After this, mice were placed back into their home cage for 1h. Following this, all mice were subjected to the first recognition phase of the OF/NOR series (OF/NOR2) where one of the objects was replaced with a different, or “novel” one. A second recognition phase of the OF/NOR series (OF/NOR3) followed, where the “novel” object from OF/NOR2 was again replaced with a second “novel” object. For both recognition phases, the total time exploring both the “familiar” and “novel” object for each mouse was quantified and preference index (PI) was calculated as a ratio between time spent exploring the novel object and the total time exploring both objects (with a minimum of 1s exploration per object). In the OF/NOR2 test (1h interval recognition phase), all groups behaved similarly, and a similar PI was obtained (NTg-Scr 0.53±0.10, n=6; NTg-Irisin 0.47±0.12, n=5; Tg-Scr 0.48± 0.09, n=6; Tg-Irisin 0.57± 0.19, n=3; data not shown). Furthermore, to provide further evidence that AAV-Irisin has the potential to counteract memory deficits, a second recognition phase (OF/NOR3) was tested 5min after OF/NOR2. Being that all mice had an additional time with the “familiar” object in OF/NOR2, in theory, mice should have increased discriminatory capabilities in the OF/NOR3 test if their memory is preserved; thus, we expected all mice to have higher PI values in the OF/NOR3 test. While there was a large variability within each group, our results showed that Tg-Scr mice had the worst performance, with their PI falling to 0.41±0.08 (Figure 3A, n=5). However, Tg-Irisin mice were able to keep the same PI as OF/NOR2 (0.56±0.19, n=4). NTg-Scr and Irisin groups also maintained a comparable level of PI (0.61±0.11, n=7, and 0.56±0.09, n=7, respectively).

**Figure 3.**
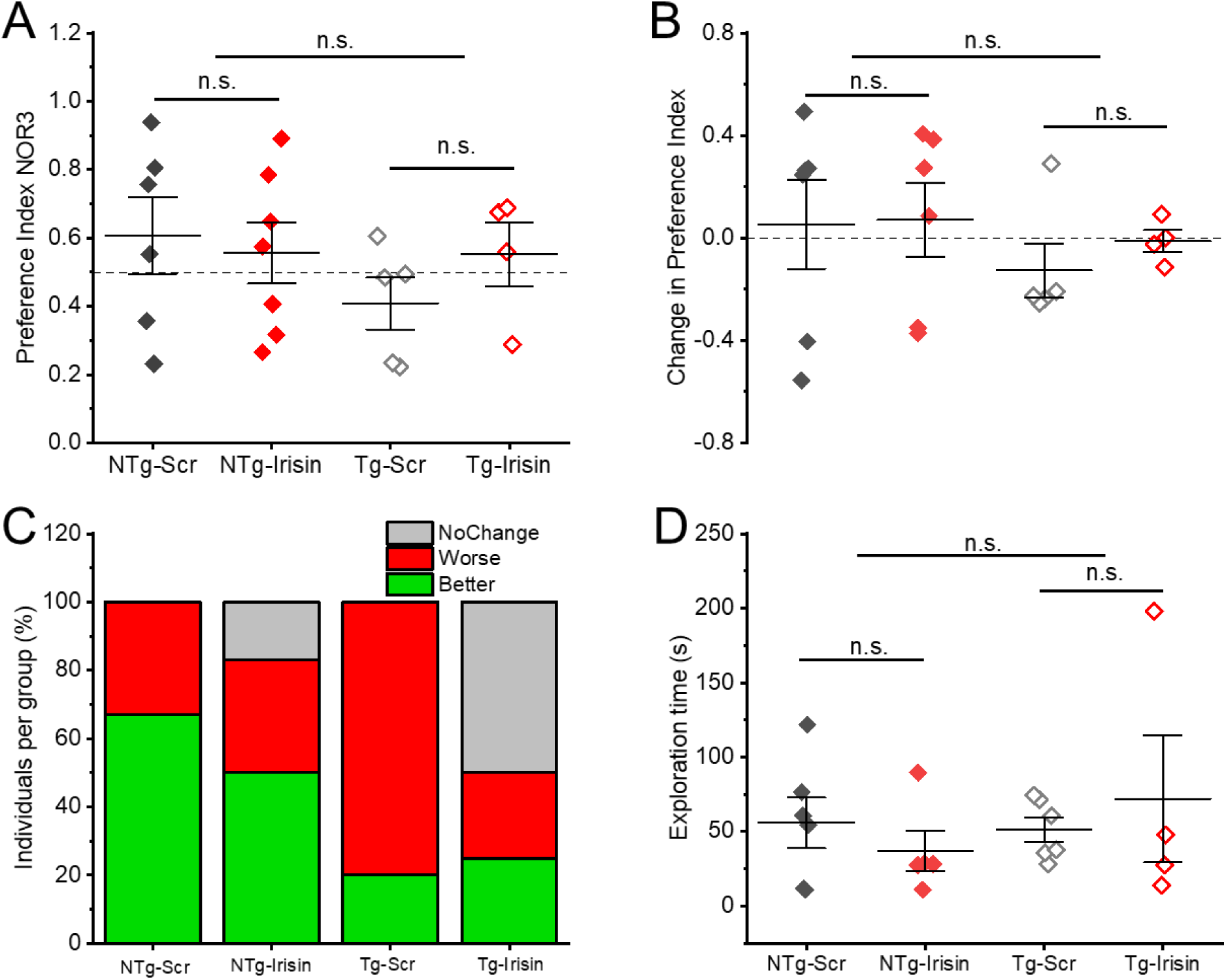
Tg-Scr mice showed the lowest performance level in the NOR test after 5 min interval (NOR3). Panel A shows preference index (PI) for the novel object amongst all groups. To better portrait these differences we calculated a new index called ΔNOR, which represents the difference between PI in NOR3 vs NOR2 (Panel B). Panel C shows the distribution of all mice within 3 categories of performance (better, worse, no change) according to ΔNOR. It is evident that most NTg mice improved in NOR3 (i.e., PI>50%) while Tg-Scr performed at the lowest level compared to other groups. Panel D shows the average total exploration time for objects in OF/NOR1 session, N.s. denotes not significantly different, p>0.05, Two-way ANOVA.

To better quantify observed differences in OF/NOR2 and OF/NOR3, we calculated a new index called ΔNOR, which was obtained subtracting PI in OF/NOR2 from PI in OF/NOR3 (Figure 3C). Improvements in OF/NOR recognition were considered if ΔNOR ≥ 0.01, and deterioration was considered if ΔNOR ≤ -0.01 (no change when -0.01 < ΔNOR < 0.01). Our results showed that most mice in NTg-Scr and NTg-Irisin groups had ΔNOR values ≥0.01 (average was 0.05±0.17 and 0.07±0.14 respectively), indicating a slight improvement in preference for novel object. On the contrary, most Tg-Scr mice showed ΔNOR<-0.01 (-0.13±0.10 on average) suggesting a reduced preference for the novel object and thus a memory deficit compared to NTg groups.

Lastly, Tg-Irisin mice showed no change on average (-0.01±0.06), indicating a slight improvement compared to Tg-Scr mice. To visualize these results and to account for variation in PI amongst individual mice within each experimental group, we segregated ΔNOR values into specific categories “better,” “worse,” or “no change” as shown in Figure 3C. Upon visual inspection, comparing within genotype, the Tg-Irisin experimental group contained 75% of mice that either improved or exhibited no change in PI in the recognition phases, compared to the Tg-Scr group that contained only 20% of mice that either improved or exhibited no change in PI in the recognition phases, and 80% of mice in that group showed worsened cognitive performance. To ensure that the observed results were not due to variations in the time exploring both objects, total object exploration from OF/NOR1 session was quantified by adding together the time exploring each individual object (Figure 3D). While there was a wide variation within groups, there was no significant difference in total time spent exploring both objects between either genotype or treatment groups. On average the total time exploring objects was 55.9±17.07s for NTg-Scr, 36.99±13.56s for NTg-Irisin, 51.34±8.09s for Tg-Scr and 71.88±42.61s for Tg-Irisin group (p=0.55 for genotype, p=0.96 for treatment).

Overall, our results show that AAV-Irisin injected directly in the hippocampus of male AD mice has the potential to reduce short-term working memory deficits. While these results show a very modest neuroprotective effect, they must be looked at in the context of AD mice whose cognitive performance is poor to begin with. It is also conceivable that muscle derived Irisin synergizes with the centrally produced myokine. Further experiments are needed to determine these differences.

### AAV-Irisin protects spatial memory in the Y-Maze Test, but May Have Anxiety-Inducing Effects

To further evaluate Irisin’s role in restoring the cognitive decline associated with AD, we performed a second behavioral test, a two-trial spatial Y-Maze test, that is known to evaluate short-term memory, just as the combined OF/NOR test does ^17,18^. During the habituation phase, mice were placed in the middle arm of the Y-maze which we labeled “Start” arm, and were only allowed to explore one of the lateral arms, designated as the “Familiar” arm for 10min. Mice were removed and placed back into their respective cages for 5min, and the blockade barricading the “Novel” arm was removed, allowing for mice to now enter this arm of the Y-Maze. Following this 5min interval mice were placed back in the same “Start” arm and were now allowed to freely explore all three arms of the Y-Maze as shown in Figure 4A (“Start,” “Familiar,” and now “Novel).

**Figure 4.**
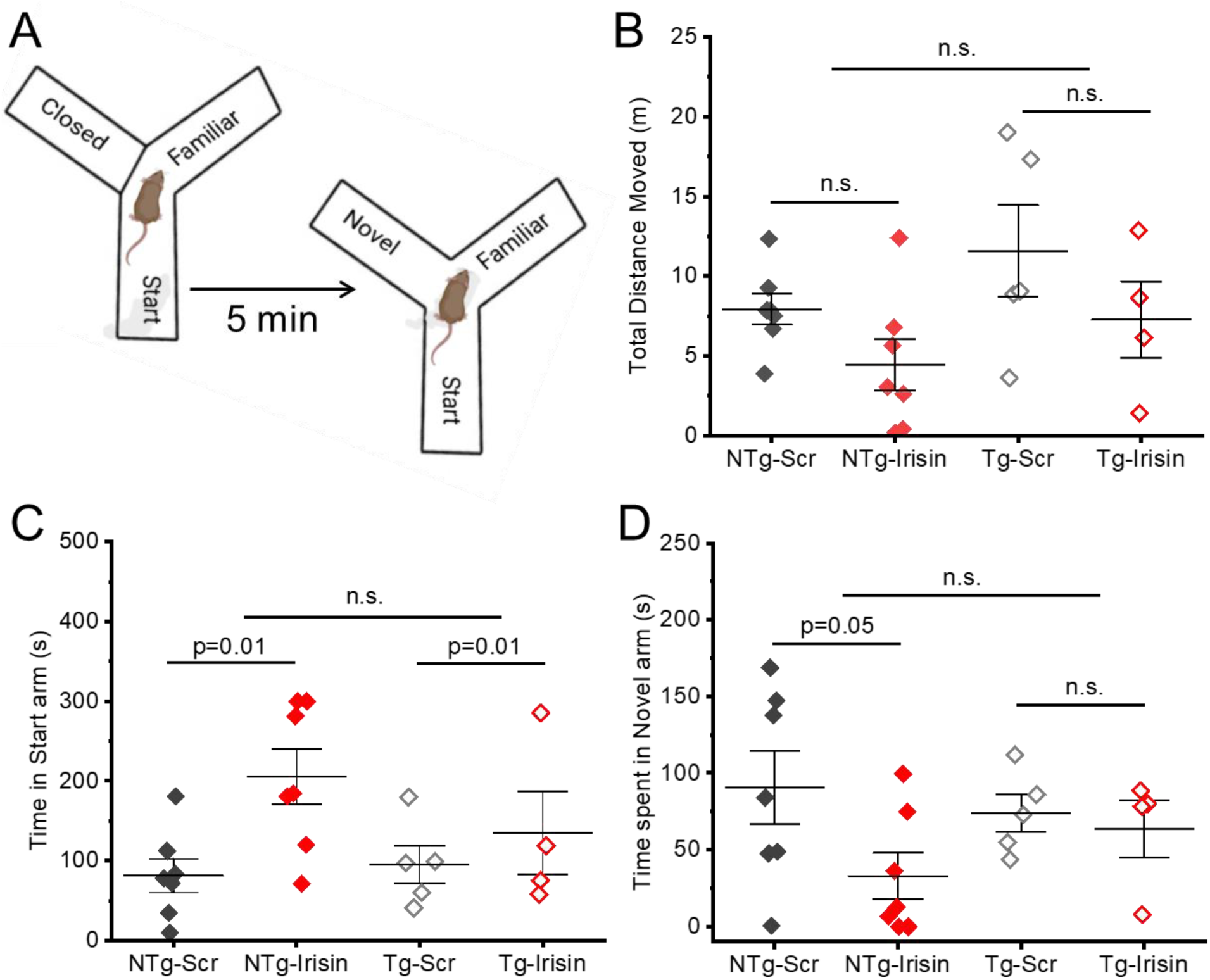
Performance in the Y maze reveals increased anxiety-like behaviors in mice injected with AAV-Irisin. (A) shows a schematic of our Y maze apparatus and experimental design. The maze on the left shows the “novel” arm being blocked during the training session, and on the right side, being open during the testing session. (B) shows the average total distance traveled during the testing session. Panel C shows the average time spent in the “start” arm of the Y maze, where mice were initially placed during the training session. Panel D shows the time spent in the “novel” arm during the testing session. N.s. denotes not significantly different, p>0.05, Two-way ANOVA.

We first assessed overall locomotor activity in the Y-Maze. We tracked the total distance moved by each mouse in the recognition phase, computing total distance moved in all three arms plus the center zone. As expected, Tg-Scr mice showed increased locomotor activity compared to NTg mice, although the difference was not as pronounced as in the combined OF/NOR test (Figure 4B), nor statistically significant between any groups. On average, Tg-Scr mice traveled the largest distance covering 11.51±1.99m (n=7) in the Y-maze, followed by Tg-Irisin mice (9.36±2.79m, n=5), NTg-Scr mice traveling 8.59±0.82m, and NTg-Irisin mice traveling 6.11± 1.76m (n=7 for both groups).

Because of the results we found in the OF/NOR testing series, we expected the same mice to show similar preference for the “Novel” arm as they did for the “Novel” object. However, when assessing the total time spent in each of the Y-Maze arms, we observed different results, especially when considering the “Start” arm (Figure 4B). In fact, mice injected with AAV-Irisin spent significantly more time in the Start arm compared to other treatments, regardless of genotype. NTg-Scr mice spent on average 81.69±20.87s in the Start arm (n=7), compared to 205.57±34.52s observed in NTg-Irisin group (p=0.01, n=7). Similarly, Tg-Scr mice spent 95.39±23.83s in the Start arm (n=5) vs 134.46±51.94s observed in Tg-Irisin mice (p=0.01, n=4). The difference between genotypes did not reach statistical significance (p=0.366). Similarly, time spent in the “novel” arm did not differ amongst groups. NTg-Scr mice spent on average 90.76±23.55s in the “Novel” arm, while NTg-Irisin group spent 32.89±15.01s (p=0.05), Tg-Scr mice spent 73.94±11.98s and Tg-Irisin mice spent 63.68±17.81s (p=0.707). These results were intriguing and suggested that injection with AAV-irisin played some role in both preserving memory function and inducing anxiolytic behaviors at the same time. A potential explanation is that mice were the most familiar with the “Start” arm of the Y-maze as they were initially placed into this arm during both the habituation and recognition phases. Thus, if mice felt threatened, stressed, or anxious, they could retreat to this “most familiar” arm where they felt the safest had their memory not decayed; injection with AAV-Irisin thus seems to play a role in both boosting short-term memory while potentially having anxiety-inducing side effects. Taken together, our results not only support the effects observed on memory function in the OF/NOR test but also show that Irisin may have adverse anxiety-inducing side effects in AD mouse models.

### Effects of central expression of Irisin on whole body and organ mass and metabolic markers

Previous studies have shown that circulating Irisin is positively correlated with energy expenditure acting as an exercise mimetic and thus has dramatic impacts on body weight and adipose tissue mass ^19,20^. Irisin has also been shown to crosstalk and regulate the function of various organs of the body, including the liver ^21,22^. We thus examined if our AAV-Irisin vector had the potential to exert such exercise mimetic effects when expressed centrally in the hippocampus. To assess the change in body weight, we weighed all mice just before stereotaxic injection of either AAV-Irisin or a scrambled AAV vector and then reassessed their body weight prior to behavioral testing (8-weeks post injection). Our results showed no significant change in body weight when comparing weight at the time of surgery vs 8 weeks later. However, these data showed a clear trend in mice injected with AAV-Irisin as their body weight changed less compared to mice injected with AAV-Scr, although not in a statistically significant manner (data not shown). NTg-Scr mice had an average gain of 4.44±10.5% body weight, NTg-Irisin mice had a lower body weight gain of 0.708 ± 2.24% (n=7), Tg-Scr mice had an average gain of 3.58 ± 4.76% body weight (n=5) and Tg-Irisin (n=5) mice had a slight loss of -0.91±3.95% body weight (genotype and treatment, p>0.05). Overall, we did observe a significant difference in body weight based on genotype only, with Tg mice weighing significantly less than NTg counterparts, regardless of AAV injection (Figure 5A). NTg-Scr mice weighed on average 49.59±0.79g and NTg-Irisin 51.26±2.71g (n=7 for both groups), versus Tg-Scr mice weighing 41.19±2.46g (n=7) and Tg-Irisin (n=5) 43.6±3.58g (p=0.003 between genotypes).

**Figure 5.**
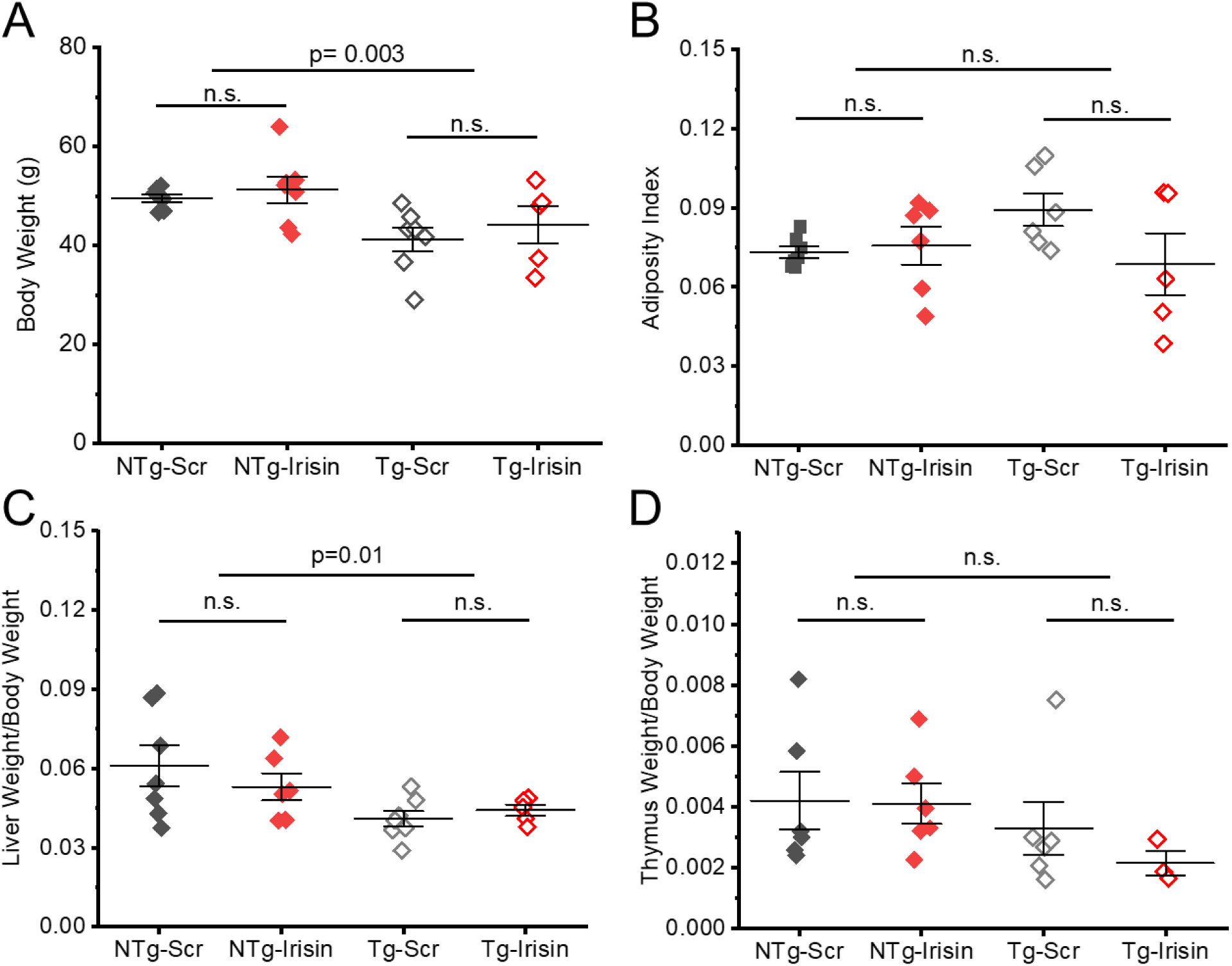
Anatomical differences between NTg and Tg mice. Panel (A) shows the average body weight at the time of behavioral analysis was conducted. Tg mice, regardless of viral injection, weighed less than age-matched NTg counterparts. (B) shows average adiposity index, calculated as a ratio between total adipose tissue and body weight. Panel C and D show the average liver and thymus size, normalized by total body weight. N.s. denotes not significantly different, p>0.05, Two-way ANOVA

Because Irisin has also been shown to decrease total fat mass (subcutaneous fat, gonadal fat, perineal fat, and mesenteric fat), we also calculated the adiposity index (=total fat mass/body weight) of each mouse prior to behavioral testing. Consistent with body weight results, there was no significant difference in adiposity between genotype or treatment (Figure 5B). On average, NTg-Scr mice had an adiposity index of 0.073±0.002 (n=7), NTg-Irisin group had an adiposity index of 0.076±0.007 (n=6), Tg-Scr had an adiposity index of 0.082±0.011 (n=6), and Tg-Irisin group (n=5) had an adiposity index of 0.069±0.012 (genotype and treatment, p>0.05). While these differences were not statistically different, it is interesting to observe that Tg-Scr mice showed a slight increase in adiposity which was reduced when injected with AAV-Irisin.

Lastly, because of Irisin’s known role in metabolic regulation and energy expenditure, we sought to understand the myokine’s relationship to liver weight. While no significant difference was found between treatments, there was a significant difference in liver weight when comparing genotypic groups (Figure 5C). To account for body weight differences, we normalized liver weight by each mouse’s own body weight. NTg-Scr mice had a normalized liver weight of 0.061±0.008 (n=7) and NTg-Irisin mice had a liver weight of 0.053±0.005 (n=6), while Tg-Scr mice had a liver weight of 0.041±0.003 (n=7) and Tg-Irisin (n=5) mice had a liver weight of 0.044±0.021 (p=0.01 between genotypes; p=0.718 between Scr and Irisin groups).

These results indicate that while AAV-Irisin may not have a direct impact on liver weight, there are genotype dependent changes that need to be further elucidated. We also investigated possible differences in thymus size, as a proxy for inflammatory status (Figure 5D). Our results showed a similar trend with smaller thymus in Tg mice, regardless of AAV injection, however this difference did not reach statistical significance. NTg-Scr mice had a normalized thymus weight of 0.0042±0.0009 (n=6) and NTg-Irisin mice had a normalized thymus weight of 0.0041±0.0007 (n=6), while Tg-Scr mice had a normalized thymus weight of 0.0033±0.0009 (n=6) and Tg-Irisin mice (n=3) had a normalized thymus weight of 0.0022±0.0004 (p=0.156 between genotypes; p=0.731 between Scr and Irisin groups).

Exercise mimetics, such as Irisin, not only increase metabolic activity, thus allowing for body weight and fat loss, but also have stress regulating effects involving anti-inflammation, anti-oxidation, and glucose homeostasis mechanisms ^23^. To get a better idea of metabolic changes associated with AAV-Irisin injection, we measured blood markers of metabolism, including glucose, ketone bodies and lactate levels in serum obtained after behavioral experiments. Figure 6 shows that blood glucose (A) and lactate (B) were comparable amongst all experimental groups, however ketone bodies levels were differentially altered in a genotype dependent manner (Figure 6C). The average blood glucose (measured in non-fasting conditions) in NTg-Scr mice was 201.75±6.38mg/dL (n=4) versus 242.0±14.95mg/dL in NTg-Irisin mice (n=6), Tg-Scr group had an average blood glucose of 236.8±10.88mg/dL (n=5) while Tg-Irisin mice (n=5) had a level of 237.6±19.16mg/dL (p=0.52 between genotypes; p=0.23 between Scr and Irisin groups). Similarly, the average lactate concentration in NTg-Scr mice was 5.32±1.44mg/dL (n=6) while in the NTg-Irisin group this was 5.46±1.22 mg/dL (n=7). The Tg-Scr group had an average blood lactate of 4.13±1.01mg/dL (n=7) compared to 3.15±1.01mg/dL in Tg-Irisin mice (n=4, p=0.19 between genotypes; p=0.96 between Scr and Irisin group). Lastly, the average level of ketone bodies in NTg-Scr mice (n=4) was 0.475±0.048 mmol/L versus 0.717±0.031mmol/L in NTg-Irisin mice (p=0.036, n=6), in the Tg-Scr group the average ketone level was 0.62±0.049mmol/L (n=5) while Tg-Irisin mice (n=5) had a level of 0.5±0.084 (p=0.39 between genotypes; p=0.38 between Scr and Irisin group).

**Figure 6.**
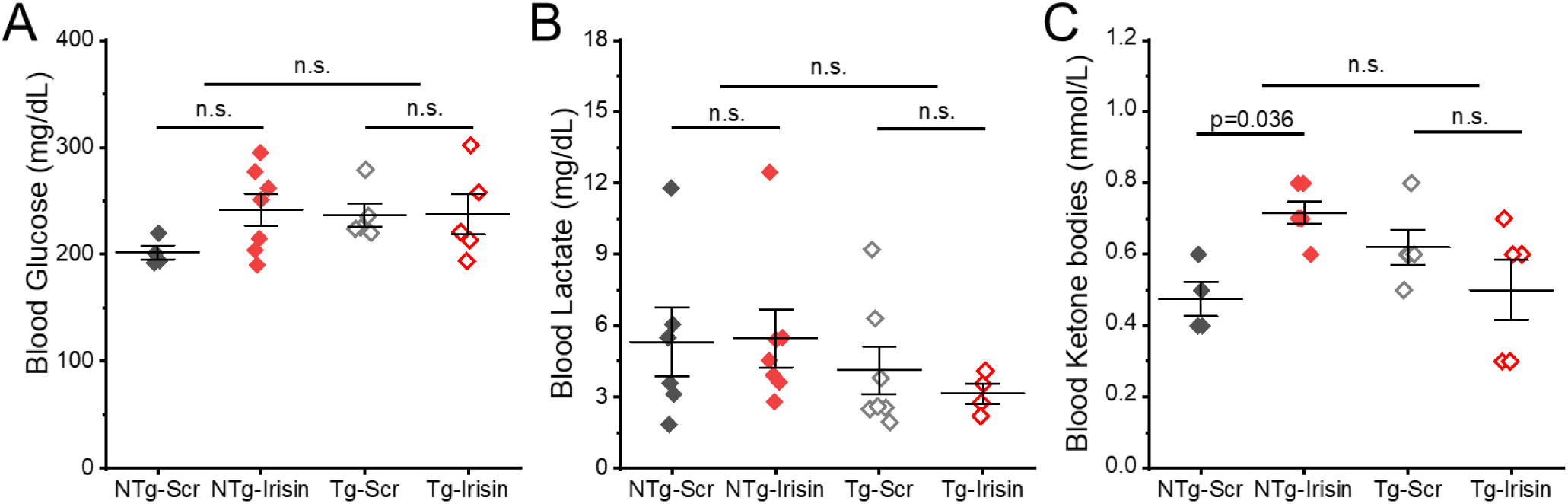
Changes in blood levels of metabolic makers including not-fasting glucose (A), lactate (B) and ketone bodies (C) at the end of all behavioral testing. Trunk blood was collected and used to immediately quantify metabolic markers. N.s. denotes not significantly different, p>0.05, Two-way ANOVA.

Overall, by assessing body and organ weight, and key metabolic markers in blood, our results show that centrally overexpressed Irisin may have slight effects compared to the peripherally released myokine. Further experiments are needed to clarify this differential effect.

## Discussion

In this study, we showed that bilateral injection of an AAV vector designed to overexpress the exerkine Irisin in the hippocampus of adult CRND8 male mice, a well-known AD mouse model, can replicate some of the effects previously observed after exercise, potentially mimicking its neuroprotective effects. While modest, our results showed that when testing short-term memory, there was a clear trend: Tg mice that were injected with AAV-Irisin did not display the same level of decline in memory function that control (Scr) mice showed in the NOR test. Interestingly, our results also showed a perhaps detrimental effect in the Y maze performance, as AAV-Irisin injected mice, regardless of genotype, spent more time in the familiar environment and less in the novel arm. One interpretation could be an increase in anxiety-like behaviors, which is also supported by the increase in grooming behavior in the OF test.

Resistance exercise, involving the use of external weights to enhance muscle strength, and endurance exercise, characterized by aerobic activities such as running and cycling, have been shown to elevate plasma and brain Irisin levels in humans, contributing to the neuroprotective effects attributed to this exerkine ^24–26^. Similar results have been observed in murine models, where regular engagement in either exercise modality led to comparable increases in plasma and brain Irisin, further supporting the role of physical activity in promoting cognitive protection ^27,28^. Despite these promising findings, practical challenges arise in applying Irisin-based interventions to neurodegenerative diseases. Aging, both physiologically and neurologically, remains the predominant time-dependent factor contributing to neurodegenerative disorders and is strongly correlated with the onset of cognitive decline ^29^. In AD, aging is associated with the accumulation of neurofibrillary tangles and β-amyloid plaques, hallmark pathological markers of disease progression ^30,31^. Additionally, aging is accompanied by physical decline, including increased sarcopenia, bone loss, reduced joint flexibility, and diminished aerobic capacity, which are exacerbated in older adults with neurodegenerative conditions ^32^. Thus, while exercise may increase Irisin levels, the physical deterioration that accompanies aging might limit the practical application of exercise-based interventions for mitigating neurodegenerative decline, particularly in AD. In the present study, we showed that non-physical activity-based overexpression of Irisin using AAV gene therapy could reduce the decline in working memory associated with AD pathology, without the need for physical exertion. Given the limited physical capabilities of the aging population and the impracticality of exercise at the intensity required to elevate Irisin levels, AAV-mediated delivery of Irisin offers a promising alternative to exercise-based therapeutic strategies for attenuating cognitive decline in AD and other neurodegenerative diseases. It is also important to point out that our results were modest, and it is possible that in combination with exercise or Irisin coming from peripheral tissue (muscle and/or adipose tissue, for example), the neuroprotective effects can be maximized. Further studies are warranted to optimize this approach and fully elucidate its potential.

AD, primarily recognized as a neurocognitive disorder that impairs memory, is frequently associated with neuropsychiatric symptoms, most notably anxiety, in humans ^33,34^. This comorbidity affects nearly 75% of AD patients, irrespective of age, indicating a strong correlation ^33^. Similarly, triple-transgenic AD mice (3xTg-AD) have exhibited significant increases in anxiety-related behaviors, as measured by standard behavioral assays such as the OF test, elevated plus maze, and light/dark box tests ^35,36^. Given Irisin’s role in mitigating the neurodegenerative effects of AD, its potential to alleviate associated comorbidities is of particular interest. Aerobic exercise, which correlates with increased brain Irisin levels, has been linked to reduced anxiety-like behaviors in mice, suggesting that locally produced Irisin may play a role in anxiety regulation ^37^. Furthermore, both long- and short-term Irisin administration has been shown to reduce anxiety-related symptoms associated with AD, highlighting a promising therapeutic strategy for addressing both the neurocognitive and neuropsychiatric symptoms of AD ^38,39^. Remarkably, the current study yielded contrasting results. While no significant differences in time spent in the center of the arena were observed between genotype and treatment groups in the OF test, an unexpected trend was noted in the Y-Maze test. Regardless of genotype, mice overexpressing Irisin spent significantly more time in the “Start” arm compared to any other arm. This result may be attributed to the familiarity of the “Start” arm, as it was the location where the mice were initially placed for behavioral testing. Interpreting this behavior, we reasoned that the mice, particularly those with Irisin overexpression, exhibited a preference for the most familiar environment, suggesting a heightened sense of comfort and/or improved memory of a familiar place. Under normal conditions, mice with intact memory are expected to explore the “Novel” arm of the Y-Maze, yet Irisin-expressing NTg mice displayed the opposite behavior, favoring the “Start” arm and spending a significantly high amount of time there. This finding suggests that while Irisin may attenuate some neurocognitive deficits associated with AD, it appears to simultaneously induce anxiety-like behaviors. The observed preference for the familiar “Start” arm aligns with typical anxious behavior in mice, where they seek environments perceived as safe and protected. While these findings contradict previous evidence supporting Irisin’s anxiolytic effects, the observed increase in anxiety should not be overlooked if Irisin is to be considered as a novel therapeutic for AD. It is also important to consider experimental differences that could result in such opposite effects. For instance, in the work of Pignataro et. al (2022 and 2023), mice were much younger than our mice (4-5 vs 12-15 months old) and Irisin treatment was administered acutely by systemic injection. In this case, the effects of Irisin could be associated to different brain regions involved in anxiety-like behaviors.

Our study also showed important systemic and metabolic differences between NTg and TgCRND8 AD mice. Tg mice were smaller in body weight, and had significantly smaller livers, even when normalized by body weight. AAV-Irisin also caused a slight reduction in adiposity index in Tg mice only (although this change did not reach statistical significance). These findings are interesting considering that dysfunctional energy metabolism, lipid metabolism, and oxidative stress are early signs of AD and related dementias neuropathology ^40^. Moreover, recent studies have shown that the liver is the first organ to experience metabolic dysregulation with progression of amyloid pathology in the amyloid precursor protein/presenilin 1 (APP/PS1) mouse line ^41^. This study showed a hypometabolic liver as early as 5 months of age in the APP/PS1 mice, with most of the disturbed metabolites being involved in several energy metabolism pathways, including amino acid and nucleic acid metabolism, as well as ketone and fatty acid metabolism. In fact, our quantification of blood metabolic markers such as glucose, lactate and ketone bodies showed that only the latter were differentially regulated in NTg mice, not in Tg counterparts. It is possible that centrally overexpressed Irisin might have reached blood circulation and directly affected liver function in NTg mice, an effect that cannot be achieved in Tg mice, due to their already dysfunctional liver. While we did not assess hepatic Irisin levels in our injected mice, more research is needed to fully characterize these changes.

